# Unsupervised content-preserving transformation for optical microscopy

**DOI:** 10.1101/848077

**Authors:** Xinyang Li, Guoxun Zhang, Hui Qiao, Feng Bao, Yue Deng, Jiamin Wu, Yangfan He, Jingping Yun, Xing Lin, Hao Xie, Haoqian Wang, Qionghai Dai

## Abstract

The development of deep learning and the open access to a substantial collection of imaging data provide a potential solution to computational image transformation, which is gradually changing the landscape of optical imaging and biomedical research. However, current implementations of deep learning usually operate in a supervised manner and the reliance on a laborious and error-prone data annotation procedure remains a barrier towards more general applicability. Here, we propose an unsupervised image transformation to facilitate the utilization of deep learning for optical microscopy, even in some cases where supervised models cannot apply. By introducing a saliency constraint, the unsupervised model, dubbed as Unsupervised content-preserving Transformation for Optical Microscopy (UTOM), can learn the mapping between two image domains without requiring paired training data and avoid the distortion of the image content. UTOM shows promising performances in a wide range of biomedical image transformation tasks, including *in silico* histological staining, fluorescence image restoration, and virtual fluorescence labeling. Quantitative evaluations elucidate that UTOM achieves stable and high-fidelity image transformations across different imaging conditions and modalities. We anticipate that our framework will encourage a paradigm shift in training neural networks and enable more applications of artificial intelligence in biomedical imaging.

---

Deep learning^1^ has made great progress in computational imaging and image interpretation^2,3^. As a data-driven methodology, deep neural networks with high model capacity can theoretically approximate arbitrary mapping from the input domain to the output domain^4,5^. Such capability offers a promising solution to image transformation, which is one of the most essential applications in biomedical imaging. The purpose of image transformation is to convert one type of image into another with highlighted or previously inaccessible information. The information extractions and predictions during transformation are beneficial to biomedical analyses by making imperceptible structures and latent patterns visible.

Recently, several network architectures have been employed for image transformation. U-Net^6^ is one of the most popular convolutional neural networks (CNNs), which has been demonstrated to have good performance in cell segmentation and detection^7^, image restoration^8^, and 3D fluorescence prediction^9^. Some elaborately designed CNNs can also complete image transformation tasks like resolution improvement^10^ and virtual fluorescence labeling^11^. Moreover, generative adversarial networks (GANs), an emerging deep learning framework based on minimax adversarial optimization that trains a generative model and an adversarial discriminative model simultaneously^12,13^, can learn a perceptual-level loss function and produce more realistic results. GANs have been verified effective in different transformation tasks such as super-resolution reconstruction^14,15^, bright-field holography^16^, and virtual histological staining^17^.

The superior performance of current deep learning algorithms largely depend on substantial high-quality training data. In conventional supervised learning, vast amounts of images and corresponding annotations are necessary, which is time-consuming and error-prone when collected manually, especially for image transformation tasks requiring pixel-level registration. Although data augmentation and transfer learning have been widely employed to reduce training data scale, collecting a small number of aligned image pairs still necessitates hardware modifications and complicated experiment procedures. In some cases, strictly registered training pairs are impossible to obtain because of the fast dynamics of biological activities or incompatibility of imaging modalities. In essence, the dilemma between the indispensability of annotated datasets and the dearth of paired training data obstructs the advancement of deep learning in biomedical imaging. Nowadays, the invention of cycle-consistent GAN (CycleGAN) makes unsupervised training of CNNs possible^18^. CycleGAN can transform images from one domain to another without paired data and exhibits comparable performance to supervised methods. This framework has been used in style transfer of natural images^19-21^ and medical image analysis^22-24^. As for optical microscopy, a few forward-looking studies have utilized CycleGAN to remove coherent noise in optical diffraction tomography^25^, and segmentation of bright-field images and X-ray computed tomography images^26^.

To advance the feasibility of unsupervised learning in biomedical applications, here we propose an Unsupervised content-preserving Transformation for Optical Microscopy (UTOM). By introducing a saliency constraint, UTOM can locate the image content and keep the saliency map almost unchanged when performing cross-domain transformations. Distortions of the image content can be avoided and semantic information can thus be well preserved for further biomedical analyses. Using this method, we implemented *in silico* histological staining of label-free human colorectal tissues with only unpaired adjacent sections as the training data. We also demonstrated UTOM’s ability on fluorescence image restoration (denoising, axial resolution restoration, and super-resolution reconstruction) and virtual fluorescence labeling to illustrate the capability and stability of UTOM.

## Results

### Principle of UTOM

The fundamental schematic of UTOM is depicted in Fig. 1a. Two image sets (A and B) are first collected to sample the source domain and the target domain. These two image domains are from different modalities (e.g. bright-field and fluorescent, low-SNR and high-SNR, anisotropic resolution and isotropic resolution etc.). No pre-aligned image pairs are required in these two image collections. Thus images in the source domain and the target domain can be acquired independently. A forward GAN and a backward GAN are trained simultaneously to learn a pair of opposite mappings between two image domains. Along with the cycle-consistency loss^18^, a saliency constraint is imposed to correct the mapping direction and avoid distortions of the image content. For each domain, a discriminator is trained to judge whether an image is generated by the generator or from the training data. When the network converges, the two GANs reach their equilibriums^27^, which means that the discriminators cannot distinguish images produced by their generators from the training data. An image could be mapped back to itself through the sequential processing of the two generators, and more importantly for biomedical images, the saliency map keeps high similarity after each transformation (Fig. 1b). For subsequent applications, pre-trained forward generators will be loaded and images never seen by the network will be fed into the model to get corresponding transformed images (Fig. 1c).

**Fig. 1.**
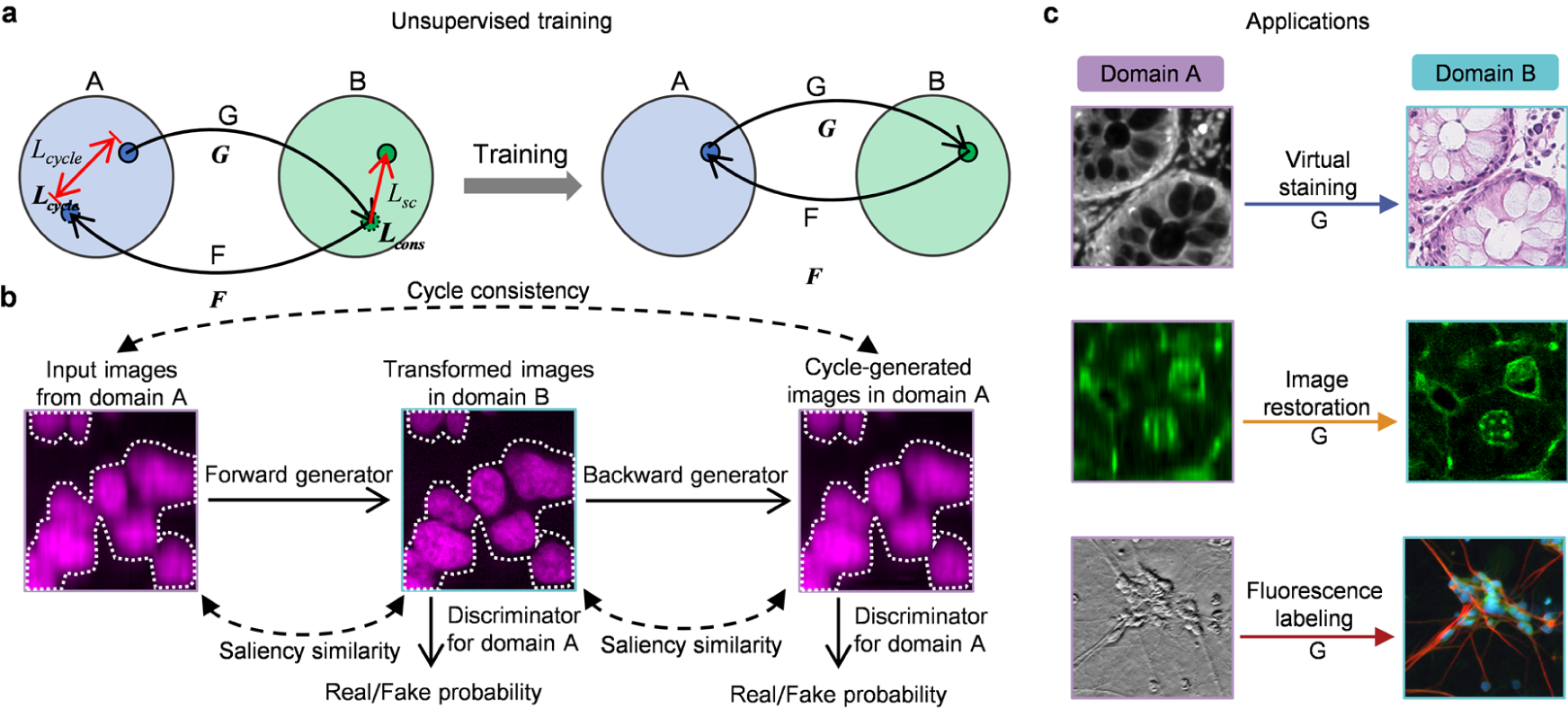
Principle of UTOM. **a**, Two sets, A and B, represent two image domains. No paired training data is required in the two sets. A forward GAN, G, and a backward GAN, F, are trained simultaneously, each for learning one direction of a pair of opposite mappings. The cycle-consistency constraint (*L*_*cycle*_) and the saliency constraint (*L*_*sc*_) are enforced to guarantee invertibility and fidelity, respectively. After proper training, the reversible mappings between the two domains can be learned and memorized in network parameters. **b**, The unfolded flowchart of UTOM. Input images from domain A are first transformed into the modality of domain B by the forward generator and then mapped back to domain A again by the backward generator. Two discriminators are simultaneously trained to evaluate the quality of the mappings by estimating the probability that the transformed images are from the training data rather than the generators. The cycle-consistency constraint is imposed to make the cycle-generated images as close to the input images as possible. The saliency constraint enforces the similarity of the image saliency (annotated by white dotted lines) before and after each mapping. **c**, Once UTOM converges, the forward generator can be loaded to perform domain transformation applications such as virtual staining, image restoration, and fluorescence labeling.

### *In silico* histological staining through unsupervised training

Firstly, we validated UTOM on transformation from label-free autofluorescence images to standard hematoxylin and eosin (H&E)-stained images. Clinically, H&E-stained histopathology slides provide rich information and are widely used for tumor diagnosis^28^. However, conventional histopathological imaging involves many complicated steps and hinders intraoperative diagnosis and fast cancer screening. Generating standard H&E-stained images from label-free imaging methods has long been pursued by biomedical researchers^29-31^. Autofluorescence images of unstained tissue can reveal histological features and the transformation from autofluorescence images to H&E-stained images is helpful for pathologists to diagnose. Conventional supervised methods for virtual histological staining^17^ rely on a laborious sample preparation and imaging procedure, which is not suitable for collecting large datasets for training clinical-grade computer-assisted diagnosis systems. UTOM can overcome these limitations by breaking the dependence on pixel-level registered autofluorescence-H&E training pairs.

We extracted samples from human colorectal tissues. After formalin fixation, paraffin embedding, and sectioning, adjacent sections were separated for the label-free processing and the H&E staining processing, respectively. We assembled tissue sections into tissue microarrays (TMAs) with tens of independent cores (see Methods). Then, we performed bright-field imaging for H&E-stained TMAs and autofluorescence imaging for label-free TMAs (Fig. 2a). In whole-slide bright-field and autofluorescence images, tissue cores in the same position are adjacent sections with similar but not identical histological features. H&E-stained adjacent sections were used as the reference to evaluate the quality of the transformation.

**Fig. 2.**
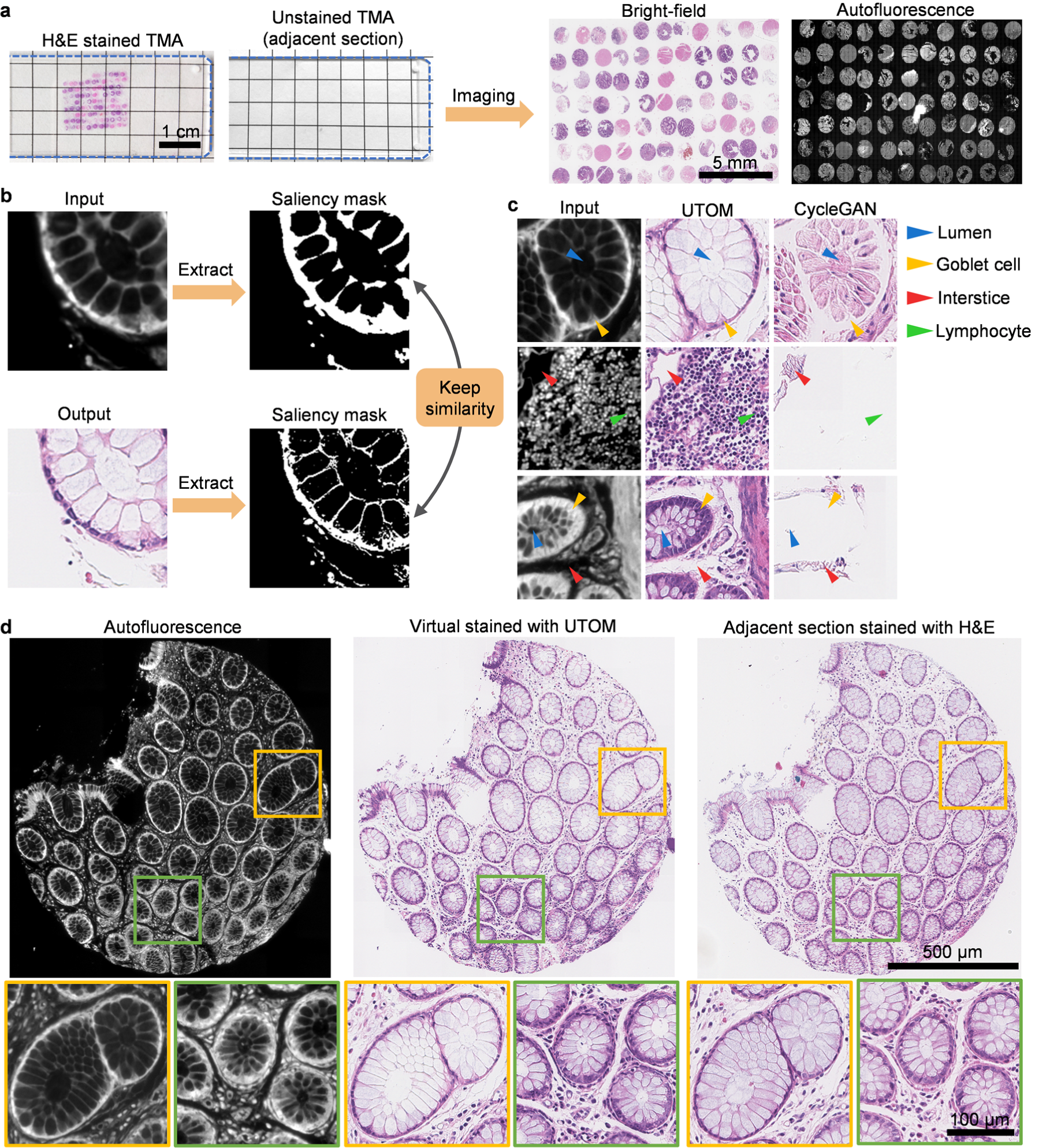
UTOM enables *in silico* histological staining of label-free colorectal slides without paired training data. **a**, Sections of colorectal tissues were assembled into tissue microarrays (TMAs). H&E staining was performed for tissue sections adjacent to the sections that skipped the staining procedure. Scale bar, 1 cm. Bright-field imaging and fluorescence imaging were performed for H&E-stained TMA and label-free TMA, respectively. Scale bar, 5 mm. **b**, Benefiting from the saliency constraint, UTOM keeps the similarity of saliency masks when images are transformed from autofluorescence to standard H&E staining. **c**, UTOM has superior convergence and stability. Histological structures can be preserved after transformation, and distortions of image contents can be avoided effectively. Arrowheads indicate different histological features including lumens (blue arrowheads), goblet cells (yellow arrowheads), interstices (red arrowheads), and lymphocytes (green arrowheads). **d**, Left: autofluorescence imaging of a label-free tissue core. Middle: H&E staining predicted by UTOM. Right: bright-field images of the corresponding H&E-stained adjacent section are shown for comparison. With UTOM, glandular architectures are well-preserved and cellular distributions are clearly resolved in a histological way. Large-image scale bar, 500 μm; enlarged-image scale bars, 100 μm.

To make the training dataset (*i*.*e*. the autofluorescence domain and the H&E stained domain), we randomly split whole-slide images of each domain into about 7000 tiles of 512×512 pixels. Small input size can reduce memory requirements and accelerate the training process. Then we trained the UTOM framework to learn the mapping from autofluorescence images to standard H&E-stained images. The saliency constraint inside UTOM can extract the saliency map of the input autofluorescence image and keep it similar when transformed to the H&E domain (Fig. 2b), which endows this framework with superior stability and reliability. Without this constraint, semantic information of the autofluorescence image is totally lost and the output H&E-stained image is distorted (Fig. 2c and Supplementary Fig. 1). This is detrimental to clinical diagnosis and could lead to severe consequences. After the content-preserving transformation of UTOM, typical histological features such as lumens, interstices, goblet cells, and lymphocytes are well preserved and become legible for pathologists. To test large images, we partition them into multiple tiles with 25% overlaps and then stitched predicted tiles together to get final results (see Methods). A tissue core virtual stained by UTOM is shown in Fig. 2d. Glandular architectures and nuclear distributions can be stained *in silico* with high fidelity and achieve comparable effect to real H&E staining. Besides visual inspection, we trained a CNN for gland segmentation (see Methods) to quantitatively evaluate whether the staining of UTOM would affect the accuracy of downstream segmentation tasks. The segmentation on UTOM-stained images achieved almost the same accuracy as on real H&E-stained images (Supplementary Fig. 2), which indicates that UTOM could accurately reconstruct H&E-stained structures from autofluorescence images and well preserve its rich histopathological information.

### Fluorescence image restoration

Next, we applied UTOM to the restoration of fluorescence images based on the experiment data released by Weigert *et al*^8^. In this series of applications, both the input and the output images are single-channel grayscale images. The target of UTOM is to learn to restore the degraded information in input images. For denoising of confocal images of Planaria^8^, we first generated the training set by randomly partitioning large images of each domain (both low-SNR and high-SNR images) into ∼18000 small 128×128 tiles (see Methods). No paired images were included in the two training sets. Then, we trained UTOM with the dataset to learn the transformation from the low-SNR domain to the high-SNR domain. Some low-SNR images that never seen by the network were used to evaluate our model and the results are shown in Fig. 3a. Our results show that UTOM can transform low-SNR images to high-SNR images while still maintaining original cell structures. We also visualized a 3D volume composed of 2856 (7×8×51) tiles to demonstrate the generalization of our model (Fig. 3b). The enhancement after denoising is remarkable and some unrecognizable details of the original noised volume become clearly visible. For quantitative analysis, we calculated image peak signal-to-noise ratios (PSNRs) before and after transformation and found that image PSNR was increased by ∼8 dB (Fig. 3c). Furthermore, we compared the performance of UTOM with 3D CARE^8^, a state-of-the-art supervised method for fluorescence image restoration (Supplementary Fig. 3a), which shows that the results of UTOM have higher fidelity and no pixel saturation occurs (Supplementary Fig. 3b). We also verified our model on restoration of multicolor zebrafish retina images with a different degradation coefficient to exhibit model capability (Supplementary Fig. 4).

**Fig. 3.**
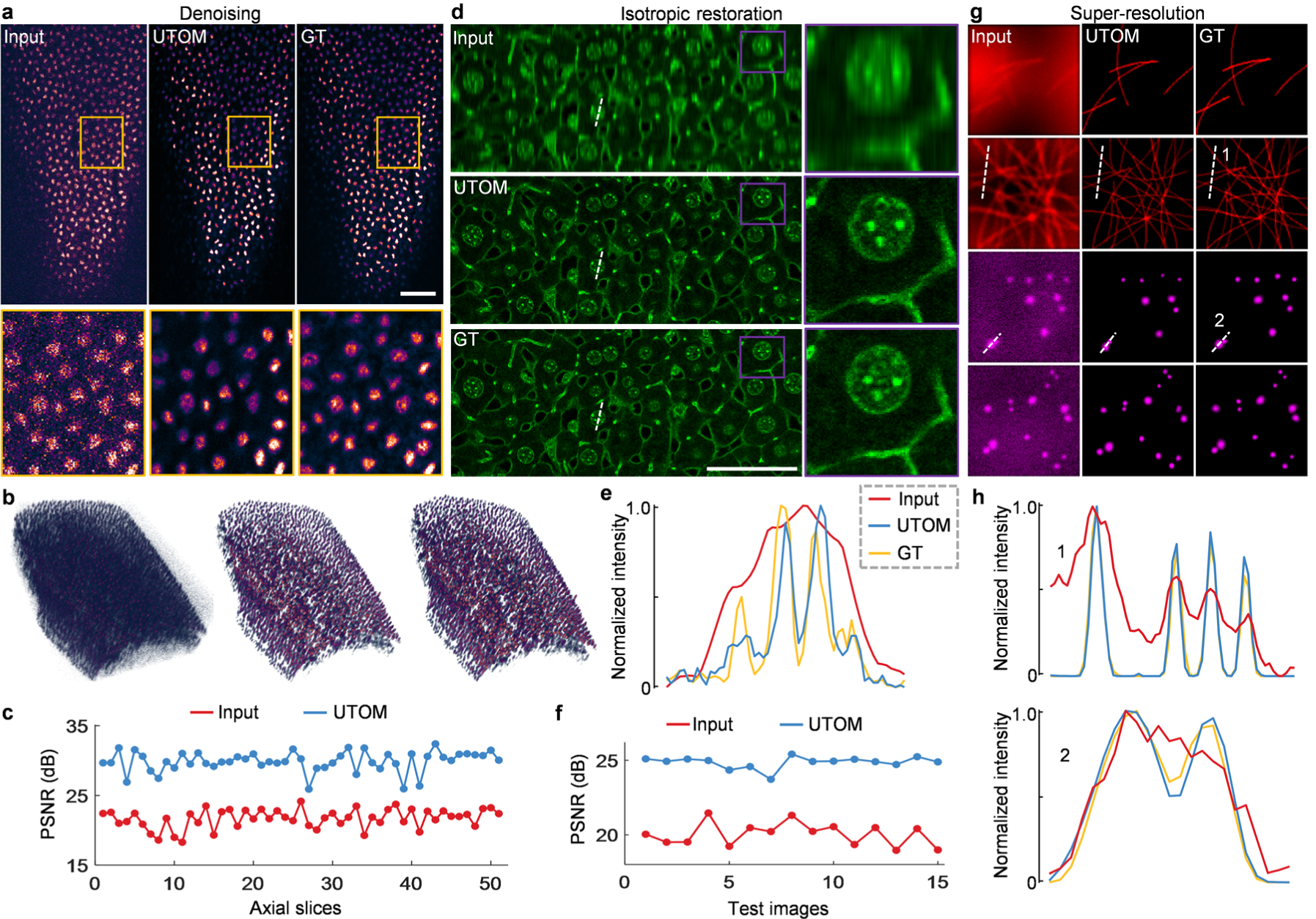
Unsupervised restoration of fluorescence images. **a**, UTOM learns to map low-SNR images to high-SNR images with high fidelity. Structures can be resolved from noised raw data imaged at low excitation dose. Scale bar, 50 μm. **b**, 3D visualization of a volume shows that some unrecognizable details become clearly visible after denoising. From left to right are the raw images, restored images with UTOM and corresponding ground truth (GT). **c**, PSNR of 51 axial slices before and after denoising, improved by ∼8 dB on average. **d**, Degradation of axial resolution can be restored by UTOM without relying on paired training data. Scale bar, 50 μm. **e**, Intensity profiles along dashed lines in **d**. Aliasing peaks in the input image can be separated by UTOM. **f**, PSNRs of 15 test images before and after isotropic restoration, improved by ∼5 dB on average. **g**, UTOM can be used to resolve sub-diffraction structures such as microtubules and secretory granules from widefield images. **h**, Intensity profiles along the dashed lines in **g**. Structures beyond recognition according to Rayleigh criterion become resolvable again.

**Fig. 4.**
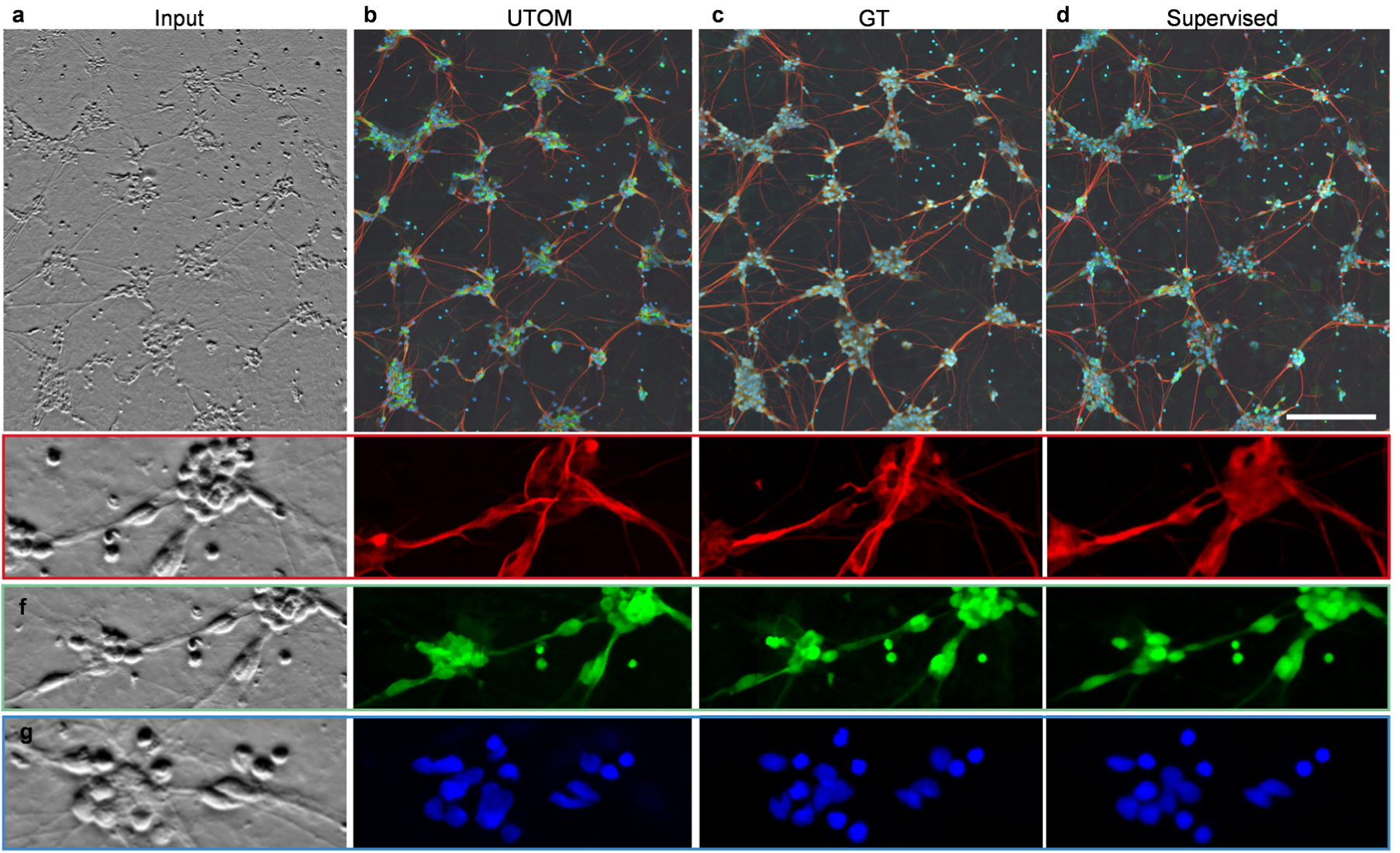
Virtual fluorescence labeling without paired training data. **a**, Phase-contrast image of differentiated human motor neurons. Original image stack is average-projected to display. **b**, Fluorescence images predicted by UTOM. **c**, Ground-truth image acquired with a spinning disc confocal microscope. Axons, dendrites and nuclei labeled by different fluorescence indicators are imaged in the red, green and blue channel, respectively. **d**, Fluorescence images predicted by the multi-scaled CNN trained with pixel-registered image pairs^11^. Scale bar, 200 μm. **e-g**, Enlargements of detailed structures extracted from the red, green and blue channel, respectively. Axons, dendrites, and nuclei can be resolved by UTOM.

Axial resolution degradation of optical microscopy can also be restored by our unsupervised learning framework. We trained UTOM with two unpaired image sets generated from the mouse liver dataset^8^ following the same procedure as denoising. As shown in Fig. 2d, an accurate mapping for axial resolution restoration was established after proper training. Blurring caused by axially elongated point spread function (PSF) was restored and some aliasing structures in original images were resolved successfully (Fig. 2e). We also quantified network performance by calculating image PSNR, as shown in Fig. 2f. After the restoration of UTOM, image PSNR was improved by ∼5 dB, average on all 15 test images. Moreover, UTOM can be used in resolving sub-diffraction structures such as microtubules and secretory granules (Fig. 2g). The transformation from wide-field microscopy to super-resolution microscopy can be learned without the supervision of any paired training data. The results of UTOM keep high accuracy and fidelity and are comparable to the CARE net^8^ (Supplementary Fig. 5). Intensity profiles of some fine structures in Fig. 2g are shown in Fig. 2f. Structures indistinguishable according to Rayleigh criterion in raw images can be recognized again with the reconstruction of UTOM.

### Virtual fluorescence labeling

Each microscopy technique has its inherent superiority and these points cannot be achieved simultaneously because of incompatible imaging principles. Our unsupervised learning framework is particularly suitable for transformations between different imaging modalities. We applied UTOM to transform phase-contrast images to fluorescence images^11^ to enable label-free imaging with component specificity. We generated an unpaired training dataset following the same procedure. Each domain contained ∼25000 image tiles. The input stack, network prediction, and corresponding ground truth are shown in Figs. 4a-4c, respectively. Most structures can be identified and labeled with correct fluorescence labels except a few neuron projections. We compared our method with conventional supervised CNN^11^ (Fig. 4d), which has a more realistic intensity distribution and better global effects. In this application, supervised learning has relatively better performance (SSIM=0.88, averaged on 3 channels) because pixel-level aligned training pairs can make the model easy to train and guide the loss function to converge to a better solution. However, UTOM is still competitive due to its independence on paired training data, which means that images of each modality can be captured at different times, in different systems, even on different samples. To evaluate each channel separately, three enlarged regions of interest (ROIs) are shown in Figs. 4e-4g and structural similarity index (SSIM) was calculated to quantify the prediction accuracy of each channel (Supplementary Fig. 6). Among the three channels, the blue channel has the best labeling accuracy with SSIM=0.87 and followed by the green channel with SSIM=0.82. The red channel has relatively large deviations with SSIM=0.64.

## Discussion

To summarize, we have demonstrated an unsupervised framework (UTOM) to implement image transformation for optical microscopy. Our framework enables content-preserving transformations from the source domain to the target domain without the supervision of any paired training data. By imposing the saliency constraint, UTOM can locate the image content and keep the saliency map similar when transformed to another domain. This improvement can effectively avoid the distortions of image content and disorders of semantic information, bringing about largely enhanced stability and reliability, which clears the obstacles for biomedical analysis. We verified this unsupervised learning framework on several image transformation tasks between different imaging conditions and modalities, such as *in silico* histological staining, fluorescence image restoration, and virtual fluorescence labeling. We evaluated the quality of the transformed images through performing downstream tasks or comparing them to corresponding ground-truth images. Quantitative metrics show that our unsupervised learning framework can learn accurate domain mappings and has comparable performance to some state-of-the-art supervised methods. Significantly, the independence on any registered image pairs during the training process makes UTOM stand out. The laborious acquisition, annotation, and pixel-level registration are no longer necessary now.

The proposed method has potential to accelerate a shift in the training paradigm of deep neural networks from conventional supervised learning to an unsupervised way. In optical microscopy, unsupervised learning has unique advantages, especially when the sample undergoes fast dynamics or preparing paired data is destructive for the sample. Moreover, combined with physical models, network parameters will be further reduced and the performance can be improved^24^. It can be expected that once the reliance on paired training data is eliminated, more applications of deep learning in optical microscopy will be made possible.

## Methods

### TMA preparation and Image acquisition

Tissues were extracted surgically or endoscopically and formalin-fixed and paraffin-embedded (FFPE). Using a tissue array instrument (Minicore Excilone, Minicore), FFPE blocks were punched and tissue cores (about 1.2 mm in diameter) were removed with a hollow needle. These tissue cores were then inserted into the recipient holes in a paraffin block and arranged into a uniform array. The re-embedded tissues were sliced into 3-μm sections. Adjacent sections were separated and mounted onto different glass slides. We performed H&E staining for one slide and kept its adjacent slide unstained. Tissue cores placed in the same position on the label-free slide and corresponding H&E-stained slide were adjacent sections for the convenience of visual inspection.

For autofluorescence imaging of label-free TMAs, we used the fluorescence imaging mode of a commercial slide scanner microscope (Axio Scan. Z1, Zeiss) equipped with a 40x/0.95 NA objective (Plan Apochromat, Nikon). We chose the DAPI excitation light source and corresponding fluorescence filter packaged in the scanner. The imaging and acquisition process (including ROI selection, field-of-view switching, autofocusing, image stitching, *etc*.) was controlled automatically by the bundled software (Zen, Zeiss). Bright-field images of H&E stained TMAs were captured with the bright-field mode of the slide scanner.

### Loss function

To ensure a reliable transformation from domain A to domain B, the mapping between the source domain and the target domain should be reversible that establishes a one-to-one correspondence. The cycle-consistency loss^18^ was used to guide the two GANs to form a closed cycle by quantifying the difference between the original images and the cycle-generated images:

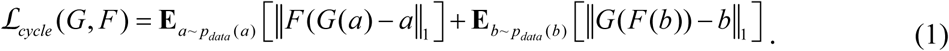

Here, **E** represents element-wise averaging and *a* and *b* are instances from domain A and domain B, respectively. *G* and *F* are the forward generator and the backward generator. The identity loss in the original cycleGAN, which is used to preserve the tint of input images, was abandoned because it was unhelpful to preserve the image content (Supplementary Fig. 7). Additionally, we imposed the saliency constraint on the loss function to realize content-preserving transformation. This constraint is based on the observation that, unlike natural scenes, the backgrounds of optical microscopy images have similar intensities. For instance, the background of fluorescence images is black while that of bright-field images is white. Using a threshold segmentation method, the rough regions of the salient objects can be extracted. Therefore, the saliency constraint is designed to be the consistency of the content masks extracted by threshold segmentation:

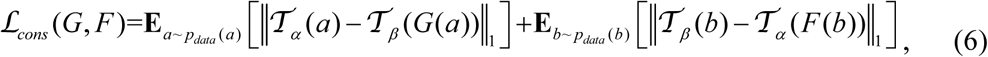

where 𝒯_*α*_ and 𝒯_*β*_ are segmentation operators parameterized by thresholds *α* and *β*. The step function of thresholding is approximated by a sigmoid function to preserve non-trivial derivatives, *i*.*e*.,

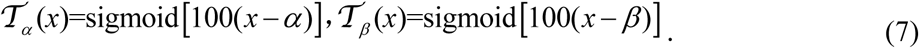

The thresholds can be automatically selected with *ImageJ*. The averaged value of all automatically selected thresholds on a subset of the training set is a good choice. We recommend checking the thresholds manually before training to ensure that the desired image content can be segmented. It is worth noting that no matter what the task is, the image content should be mapped to 1 while the background should be mapped to 0. For virtual histopathological staining, pixel intensity=0 means the background in domain A while pixel intensity=255 means the background in domain B. The segmentation operator of domain B was adjusted as

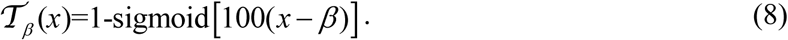

More details and some real-data examples are shown in Supplementary Fig. 8. Finally, the full loss function can be formulated as:

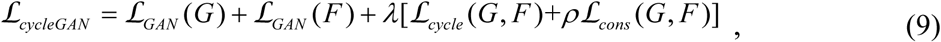

Where ℒ_*GAN*_ (*G*) and ℒ_*GAN*_ (*F*) are the adversarial losses of the forward GAN and the backward GAN, respectively (Supplementary Notes). *λ* and *ρ* are hand-tuned weights to adjust the strength of the cycle-consistency loss and the saliency constraint. Inspired by the adaptive learning rate in modern gradient descent optimizers, *ρ* is designed to be exponentially decayed during the training progress. The saliency constraint is only dominant at the beginning of the training process to guide the network to converge in a better direction. In the later stage of training, the cycle-consistency will not be weakened. The influence of *ρ* is investigated in Supplementary Fig. 9.

### Network architectures and training details

The generator and the discriminator of UTOM adopted the classical architectures of CycleGAN (Supplementary Notes and Supplementary Fig. 10). Considering the diversity of microscopy images, many image transformation tasks would change the channel number of the image and thus the two GANs must have matched input and output channel numbers. For the generator, the kernel size of the first convolution layer was set to be 7 to endow this layer with large receptive fields for extracting more neighborhood information. We used padded convolution layers to keep the image size unchanged when passing convolution layers. All the downsampling and upsampling layers were implemented by strided convolution with trainable parameters. For the discriminator, we adopted the PatchGAN^32,33^ classifier. This architecture penalizes structures at the scale of local patches to encourage high high-frequency details. More specifically, it divides the input image into small patches, and classification is performed at the patch scale. The ultimate output is equivalent to the average of the classification loss on all patches.

Adam optimizer^34^ was used to optimize network parameters. The exponential decay rates for the 1st and 2nd moment estimates are set to be 0.5 and 0.999, respectively. We used batch size of 1, and trained our network for ∼10k iterations. We used graphics processing units (GPUs) to accelerate computation. On a single NVIDIA GTX 1080Ti GPU (11GB memory), the whole training process of a typical task (single-channel images, 128×128 pixels, 50k iterations) takes approximately 10 hours. Considering the possibility of using multiple GPUs for parallel computation, the training time can be further reduced.

### Data pre- and post-processing

Histological images (autofluorescence and H&E-stained) captured with the commercial slide scanner were first converted to 8 bit tiff files using a Matlab (MathWorks) script. Tissue cores were cropped out separately from whole-slide images and divided into a training set and a test set. Then large images were randomly split into 512×512 tiles for training. For pre-processing of fluorescence images, each dataset with ground truth was divided into the training part and the test part in a ratio of about 5: 1. For each image pair in the training part, we randomly decided whether to pick the original image or its ground truth with a probability of 0.5 and discard the other one. Then, the selected images were collected into domain A and domain B, respectively. This operation guaranteed that there were no aligned image pairs in these two image sets (Supplementary Fig. 11).

In the test phase, all original images in the test set were fed into the pre-trained forward generator. Large images were partitioned into small tiles with 25% overlaps. For post-processing of transformed images, we cropped away the boundaries (half of the overlaps) of the output tiles. Stitching was performed by appending the remaining tiles right next to each other (Supplementary Fig. 12). 3D visualization was performed with the built-in *3D viewer* plugin of *ImageJ*. Reconstructed data volumes were adjusted to the same perspective for the convenience of observation and comparison.

### Gland segmentation

To reduce the amount of data required for training, the segmentation of colorectal glands was based on transfer learning. We first pre-trained a standard U-net model with the GlaS dataset^35,36^ to learn to extract generic features of colorectal glands. But we found significant differences between the tint of our H&E-stained slides and those in the GlaS dataset. To improve segmentation accuracy, we manually annotated a few images and then fine-tuned the model with our dataset. Data augmentation was adopted in both the pre-training phase and the fine-tuning phase to reduce data requirements. After training, large images were partitioned into 512×512 tiles with 25% overlaps and then fed into the pre-trained model. The predicted tiles were stitched together to form the final results. For quantitative analysis, the intersection-over-union (IoU) score between the network-segmented mask and corresponding manually annotated mask was computed^37^.

### Benchmarks and evaluations

We compared our unsupervised framework to the state-of-the-art supervised CNNs to demonstrate its competitive performance. We used the original CARE net^8^ for the task of fluorescence image restoration. For Planaria denoising, we adopted the released model embedded in the *CSBDeep* plugin of *Fiji*. This model was trained in 3D before released. For isotropic restoration and super-resolution reconstruction, we retrained the CARE net using a training set of similar size to UTOM. All images were organized in 2D in these two tasks. In the experiment of virtual fluorescence labeling, the elaborated-designed multi-scale CNN trained with pixel-level aligned image pairs^11^ was loaded and test images were fed into and flowed through the network. The released models are reliable enough to produce best results that the supervised network can achieve.

The evaluation strategy was on the basis of not only revealing the differences visually but also providing quantitative analyses of transformation deviations. The quantitative metrics we used for image-level evaluation are PSNR and SSIM. SSIM is suitable to measure high-level structural errors while PSNR is more sensitive to absolute errors at pixel level. For RGB images, SSIMs are averaged on three channels. In the task of denoising, we also calculated the histograms of pixel values using *ImageJ* to show the distributions of image intensity. For better visualizing the transformation deviations, we assigned network output and corresponding ground truth to different color channels (*i*.*e*., network output to the magenta channel, and ground truth to the green channel). Considering that magenta and green are complementary colors, if the structures before and after transformation are in the same position and with the same pixel values, these structures in the merged image will be shown in white. Otherwise, these pixels would be displayed as the color with larger pixel values. This strategy was used to independently visualize the transformation error (especially position offset) across different channels in virtual fluorescence labeling.

## Data availability

The dataset for histological staining is available from the corresponding author upon request. The data for fluorescence image restoration are available at https://publications.mpi-cbg.de/publications-sites/7207/. The dataset used during virtual fluorescence labeling can be found at https://github.com/google/in-silico-labeling/blob/master/data.md. Some data processed by our preprocessing pipeline that can be used for training and test is made publicly available at https://drive.google.com/drive/folders/1QPlLcTHlU58xo116KB1bd680EoMof_Wn?usp=sharing.

## Code availability

Our Pytorch implementation of UTOM is available at https://github.com/Xinyang-Li/UTOM. The scripts for data pre- and post-processing were also uploaded to this repository.

## Acknowledgements

We would like to acknowledge Weigert *et al*. for opening their source code and data on image restoration to the community. We thank the Rubin Lab at Harvard, the Finkbeiner Lab at Gladstone, and Google Accelerated Science for releasing their datasets on virtual cell staining. We thank Jingjing Wang at the apparatus sharing platform of Tsinghua University for the help on imaging of histopathology slides.

## Author Contributions

Q. D., H. W. and XY. L. conceived the project. Q. D. and H. W. supervised the research. XY. L., G. Z. and H. Q. designed detailed implementations, carried out experiments and tested the codes. XY. L., G. Z., H. Q., F. B., J. W. and Y. D. discussed and designed the experimental schemes. Y. H. and J. Y provided the histological slides and corresponding bright-field images. H. Q., F. B., J. W., X. L., H. X. and Y. D. gave critical discussions on the results and evaluation methods. All authors participated in the writing of the paper.

## Materials & Correspondence

Correspondence and requests for materials should be addressed to Q. D. or H. W.

