## Supplementary Information for "Unsupervised content-preserving transformation for optical microscopy"

### Supplementary Figures

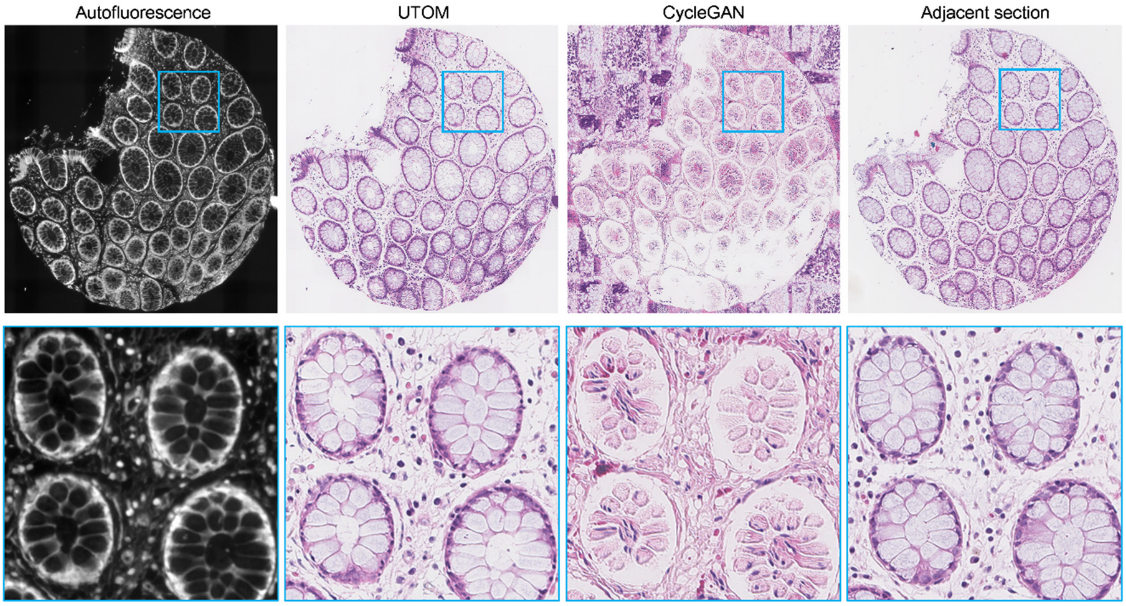


**Supplementary Figure 1 | The saliency constraint can preserve the image content during transformation.** Original cycleGAN does not have the content-preserving ability and the image content is totally distorted when transformed to the target domain. With the saliency constraint, UTOM can perform content-preserving transformations and the semantic information can be well preserved. The adjacent section stained with H&E is provided in the rightmost panel for reference.

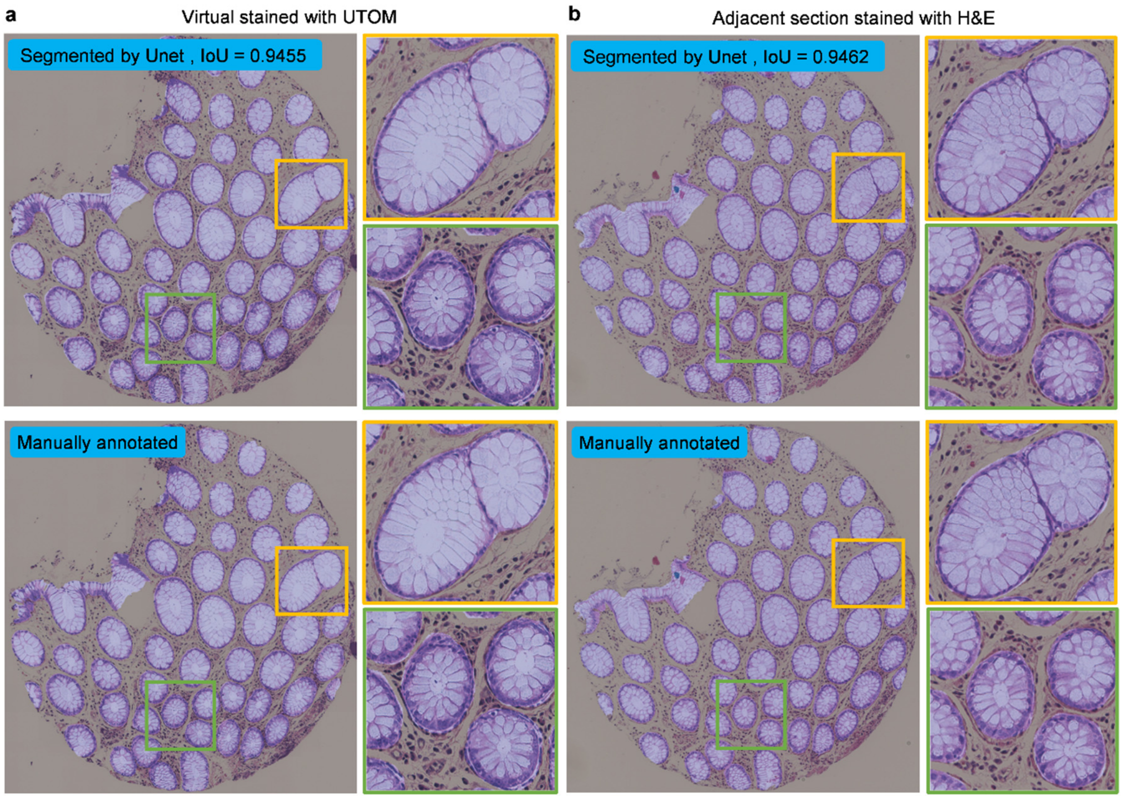


**Supplementary Figure 2 | Gland segmentation of virtual stained and H&E-stained slides.** A U-net was trained to segment secretion glands form histological images. **a**, Segmentation result of virtual stained slide (IoU=0.9455). **b,** Segmentation result of corresponding adjacent section stained with H&E (IoU=0.9462). Segmented regions were highlighted in bright purple. Manually annotated masks (bottom panel) serves as the ground truth.

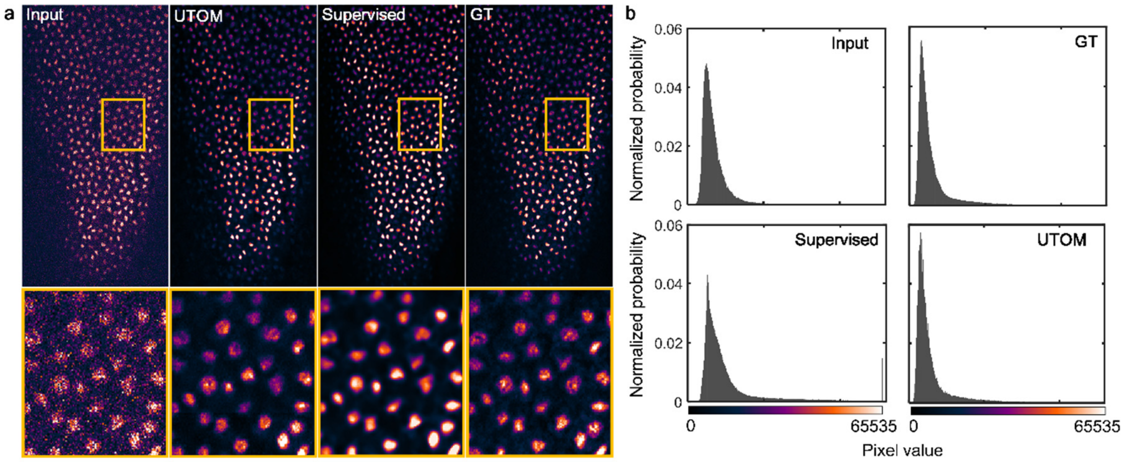


**Supplementary Figure 3 | High-fidelity denoising with UTOM. a**, The result of UTOM, the ground truth, and the result of supervised CARE net are provided for comparison. **b**, Histograms of the results reveal the distributions of pixel intensity. Overexposure is obvious in the red dashed box where pixel value is 65535. The histogram of our result is very close to the ground truth.

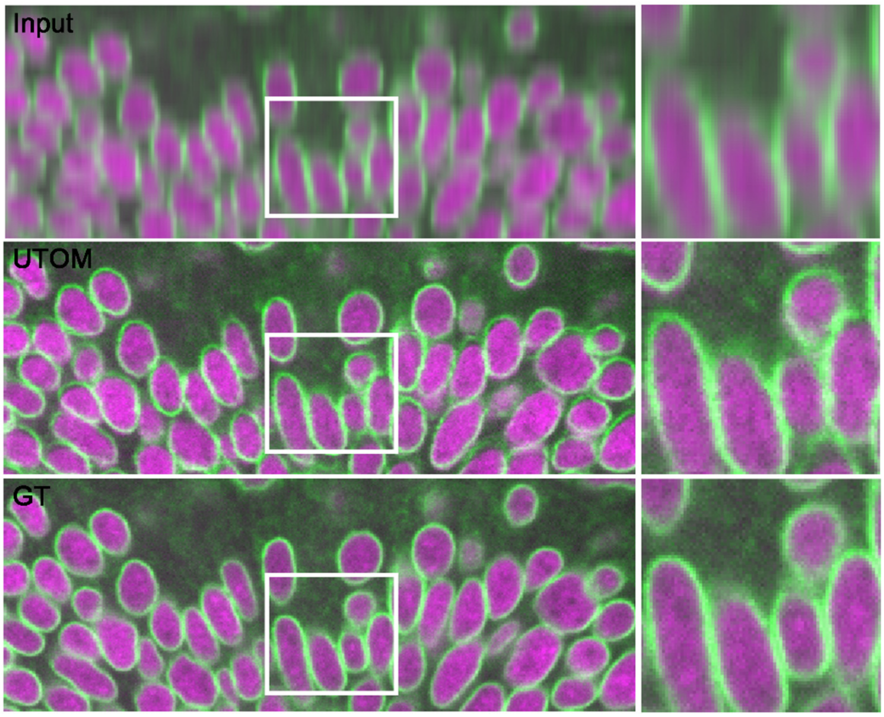


**Supplementary Figure 4 | Isotropic restoration of degraded axial resolution on zebrafish retina.** Nuclei and nuclear envelopes are labeled with DRAQ5 (magenta) and GFP-LAP2b (green), respectively. Our method can restore the resolution of the two channels to an identical level.

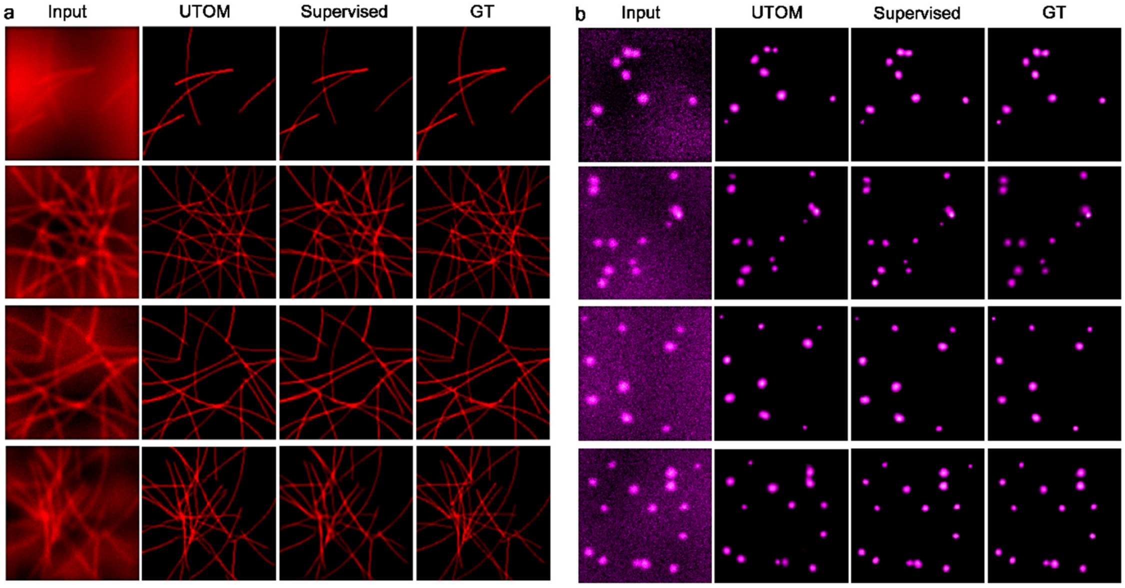


**Supplementary Figure 5 | High-fidelity super-resolution reconstruction by UTOM.** Sub-diffraction structures like **a**, microtubules and **b**, granules can be resolved from wide-field images. The original input images, the results of UTOM, the results of 2D CARE, and corresponding ground-truth images are shown in each column.

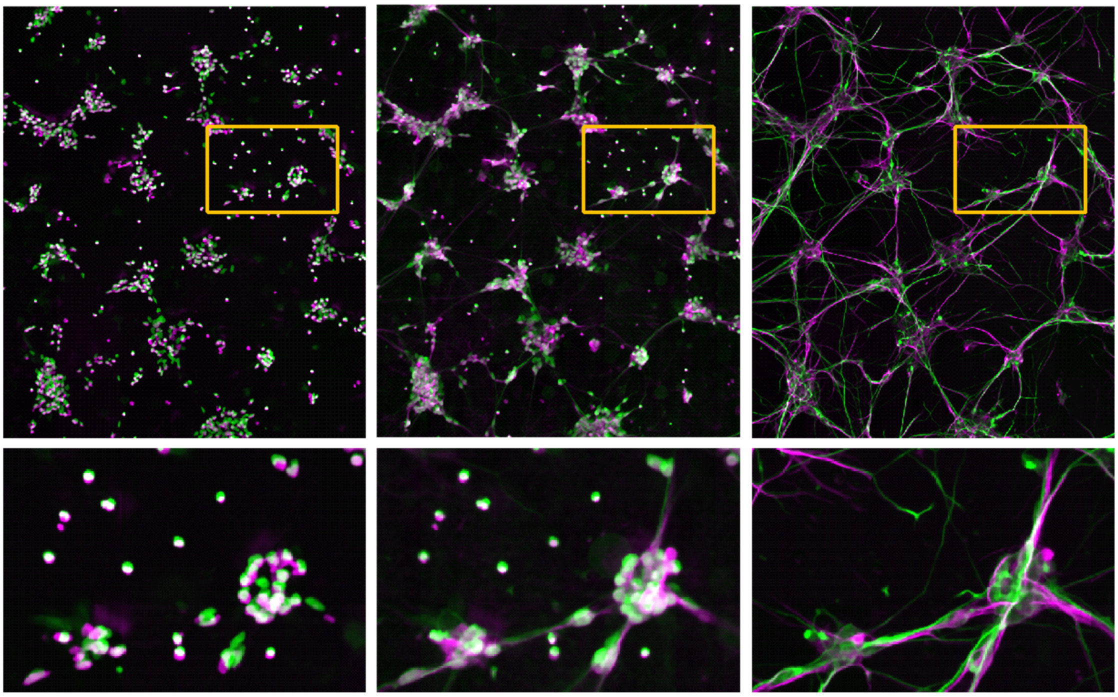

**Supplementary Figure 6 | Merged maps of virtual fluorescence labeling.** The results of UTOM (magenta) and the ground-truth images (green) are merged together. **a**, The blue channel labels nuclei of human motor neurons with SSIM=0.87. **b**, The green channel labels dendrites with SSIM=0.83. **c**, The red channel labels axons with SSIM=0.64. Images in the bottom row are enlargements of the same region.

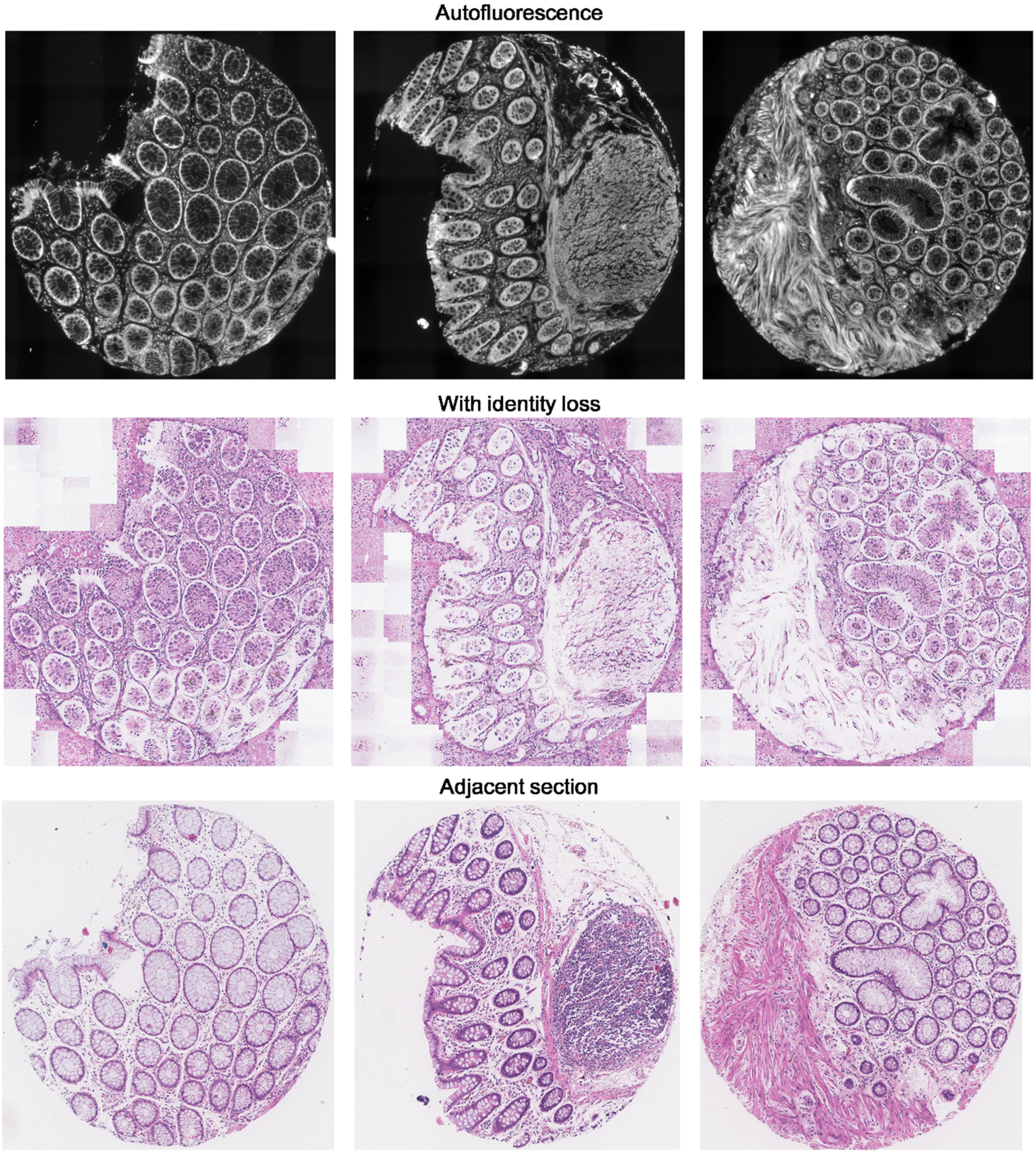


**Supplementary Figure 7 | The identity loss cannot preserve the image content in histological staining.** From top to bottom are autofluorescence images, transformed images with the identity loss, and corresponding H&E-stained adjacent sections, respectively. Three tissue cores are shown here for instance. Similar results were achieved on other tissue cores and other independent trials (N=5).

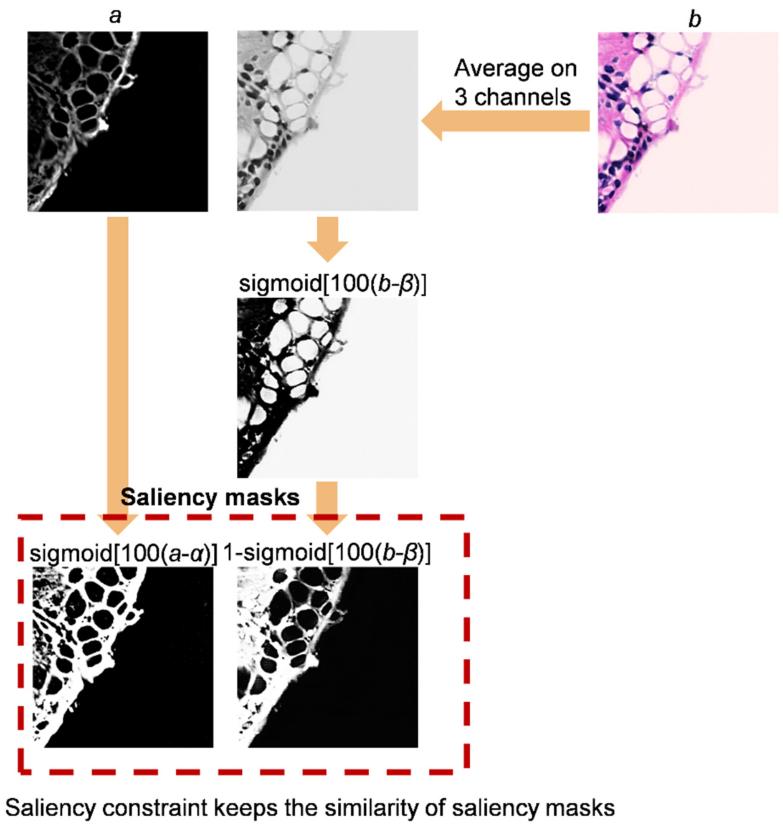


**Supplementary Figure 8 | Saliency constraint in virtual histopathological staining.** For domain A (autofluorescence images), saliency masks were extracted directly by sigmoid[100($a$-$\alpha$)]. For domain B (H&E images), we first converted the RGB images to its grayscale version by averaging the three channels. Then, the saliency masks were extracted by 1-sigmoid[100($b$-$\beta$)]. The image content should be mapped to 1 while the background should be mapped to 0.

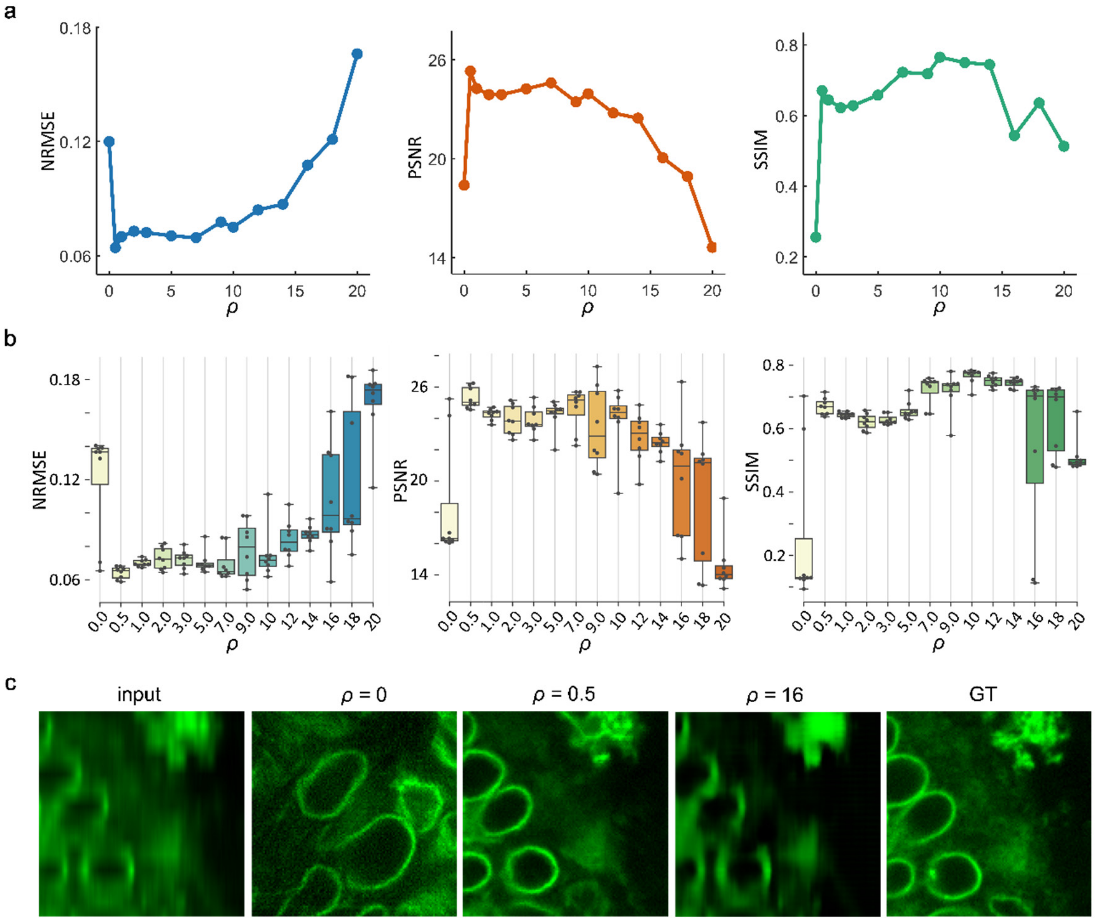


**Supplementary Figure 9 | Performance of UTOM when imposed the saliency constraint with different constant weights. a**, Averaged NRMSE/PSNR/SSIM of 8 independent experiments (each ρ) on the zebrafish retina dataset (Supplementary Fig. 4). The saliency constraint was imposed with different constant weights ranging from 0 to 20. For each experiment, NRMSE/PSNR/SSIM were arithmetically averaged on all 144 image patches in the test set. **b**, Box-dot plots show the distribution of NRMSE/PSNR/SSIM obtained with different ρ. **c**, Typical results under different ρ. Without saliency constraint, the network is unstable and often converges to wrong mappings. The saliency constraint can correct the mapping bias effectively. However, when ρ is relatively large, other terms will be less important and the performance will degrade.

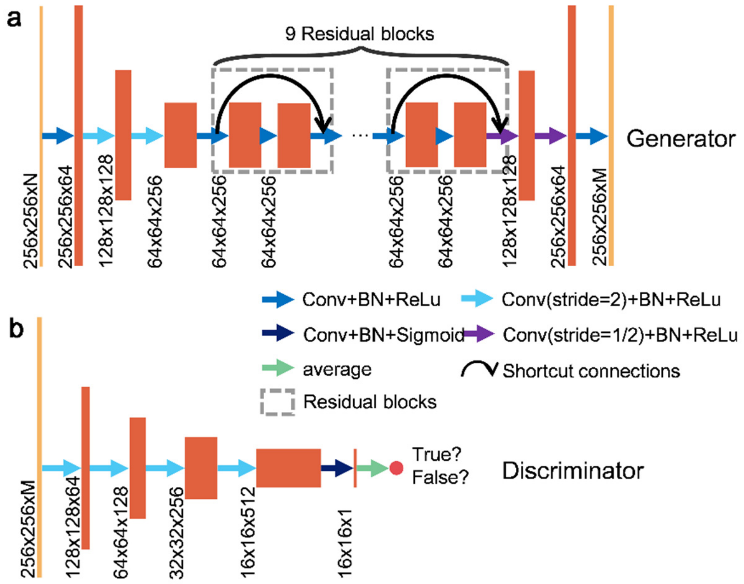


**Supplementary Figure 10 | Network architectures.** Here we take 256×256 input size as an example. Each coral rectangle represents a feature map extracted by convolutional kernels. **a**, The generator is a multi-layer residual network with downsampling input layers and upsampling output layers. **b**, The discriminator (PatchGAN classifier) uses multiple stride convolution layers for abstract representation. It first generates a matrix, where each element corresponds to a patch in the input image. The ultimate output is the average of the cross-entropy loss on all patches.

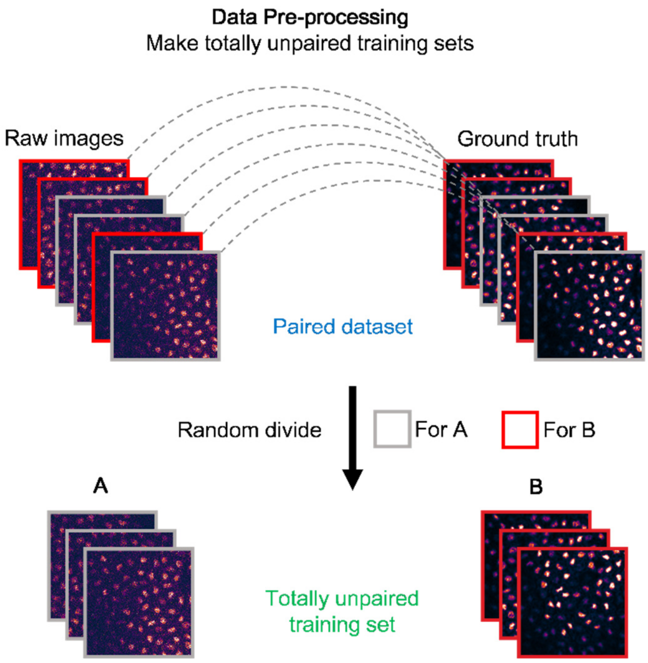


**Supplementary Figure 11 | Data pre-processing pipeline.** Most training sets were randomly selected from published datasets with paired ground-truth images. Specially, we randomly selected one half of the dataset and collected its raw images into domain A, and then selected the other half of the dataset and collected its ground-truth images into domain B.

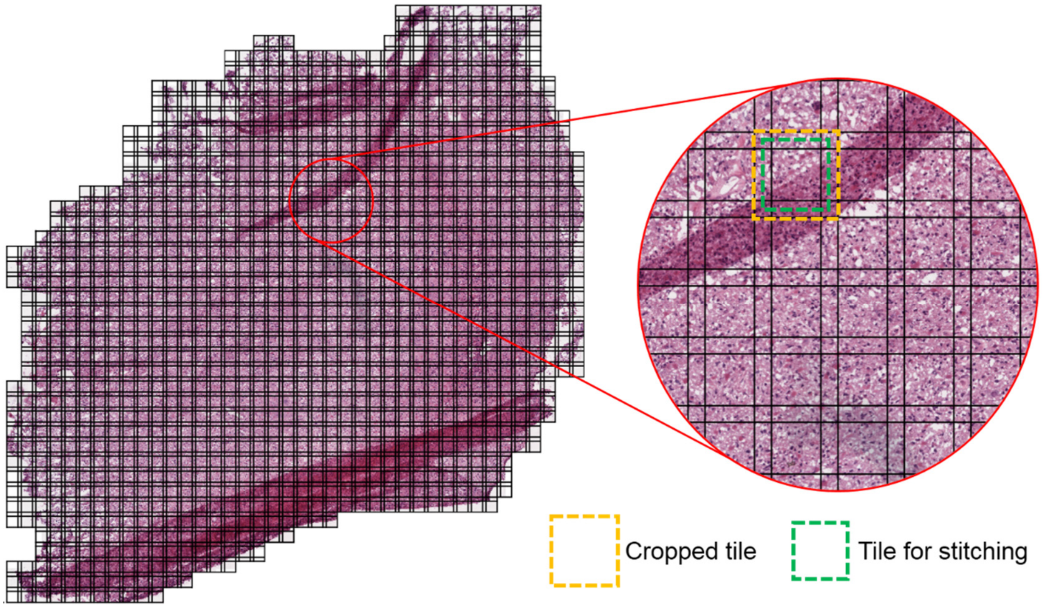


**Supplementary Figure 12 | Tiling and stitching in pre- and post-processing.** In most cases, images needed to be transformed are extremely large in pixel size. In our data processing pipeline, large images were partitioned into multiple overlapping tiles to reduce memory requirements and improve training efficiency. The edges of output patches were cut out and the rest parts were stitched together into a large image.

#### Supplementary Notes

##### Network architectures and the loss function.

UTOM is composed of two GANs to learn the mapping function between two image domains in an unsupervised way. The architectures of the generator and the discriminator are visualized in Supplementary Figs. 10. The first three layers of the generator are downsampling layers implemented by strided convolution to extract low-level abstract representations. Nine stacked residual blocks are followed to extract high-level features. The number of the residual blocks reflects model capacity. More residual blocks are recommended for more complex tasks. The last three upsampling layers are also implemented by strided convolution. They are used to integrate the features and rescale image to its original size. The discriminator is a CNN with fewer convolution layers. Each layer downsamples the feature maps but doubles the channel number. The last convolution layer extracts a single-channel feature map and classification is performed on each element of this feature map (PatchGAN classifier). The final true or false label is generated by averaging the individual labels of each element. Each convolution layer in both the generator and the discriminator contains a nonlinear activation unit. Whether to use the sigmoid function or rectified linear unit (ReLU) is indicated by corresponding arrows in Supplementary Figs. 10.

It should be noted that the input and output channel numbers of the two GANs should match to ensure to form a complete cycle, especially when the images in domain A and domain B have different channels numbers. In terms of the objective function, the first part is the frequently-used adversarial losses of the two GANs, which can be formulated as the most common form:

$$\mathcal{L}_{GAN}(G)=\mathbf{E}_{b\sim p_{data}(b)}\left[\log D_B(b)\right]+\mathbf{E}_{a\sim p_{data}(a)}\left[\log(1-D_B(G(a))\right]$$
$$\mathcal{L}_{GAN}(F)=\mathbf{E}_{a\sim p_{data}(a)}\left[\log D_A(a)\right]+\mathbf{E}_{b\sim p_{data}(b)}\left[\log(1-D_A(G(b))\right], \quad \text{(S1)}$$

where $D_A$ and $D_B$ represent the discriminator of the forward GAN and the backward GAN, respectively. Lowercase letters *a* and *b* are images from the image domains represented by corresponding capitals. $\mathbf{E}$ is the expectation operator. The second part of the loss function is the cycle-consistency loss, which is most essential for training the two GANs. The last part is the saliency constraint term to correct mapping errors and improve the success rate of training. The full objective function can be formulated as

$$\begin{aligned}\mathcal{L}_{cycleGAN} = {} & \mathbf{E}_{b\sim p_{data}(b)}\left[\log D_B(b)\right]+\mathbf{E}_{a\sim p_{data}(a)}\left[\log(1-D_B(G(a))\right]+ \\ & \mathbf{E}_{a\sim p_{data}(a)}\left[\log D_A(a)\right]+\mathbf{E}_{b\sim p_{data}(b)}\left[\log(1-D_A(G(b))\right]+ \\ & \lambda\left\{\mathbf{E}_{a\sim p_{data}(a)}\left[\left\|F(G(a))-a\right\|_1\right]+\mathbf{E}_{b\sim p_{data}(b)}\left[\left\|G(F(b))-b\right\|_1\right]\right\}+ \\ & \rho\left\{\mathbf{E}_{a\sim p_{data}(a)}\left[\left\|\mathcal{T}_\alpha(a)-\mathcal{T}_\beta(G(a))\right\|_1\right]+\mathbf{E}_{b\sim p_{data}(b)}\left[\left\|\mathcal{T}_\beta(b)-\mathcal{T}_\alpha(F(b))\right\|_1\right]\right\}\end{aligned}, \quad \text{(S2)}$$

where $\lambda$ is a constant to enforce cycle-consistency loss. $\rho$ can be constant or exponentially decayed to enforce the saliency constraint. If $\rho$ is set to be a constant, the convergence will be faster. If it is exponentially decayed, the final effect will be better because the convergence direction is only constrained at the beginning of training. Cycle consistency will not be weakened at the end of training. $\mathcal{T}_\alpha$ and $\mathcal{T}_\beta$ are segmentation operators parameterized by threshold $\alpha$ and $\beta$. They are used to extract saliency masks of the images in domain A and domain B, respectively. Here, we used sigmoid(100$x$) to approximate the Heaviside step function of threshold segmentation to keep nontrivial gradient, *i.e.*,

$$\mathcal{T}_{\alpha}(x)\text{=sigmoid}\left[100(x-\alpha)\right], \mathcal{T}_{\beta}(x)\text{=sigmoid}\left[100(x-\beta)\right]. \tag{S3}$$

It is worth noting that no matter what the task is, the image content should be mapped to 1 while the background should be mapped to 0. For virtual histopathological staining, pixel intensity=0 means the background in domain A while pixel intensity=255 means the background in domain B. The segmentation operator of domain B should be adjusted as

$$\mathcal{T}_{\beta}(x)\text{=1-sigmoid}\left[100(x-\beta)\right]. \tag{S3}$$

More details and some real-data examples are shown in Supplementary Fig. 8.
